# Protection against reinfection with *Mycobacterium tuberculosis* extends across heterologous Mtb lineages

**DOI:** 10.1101/2025.07.08.663727

**Authors:** Andrew W Simonson, Michael C Chao, Luke E Hood, Rachel A Donlan, Forrest Hopkins, Michael R Chase, Andrew J Vickers, Alanna Callendrello, Edwin Klein, H Jacob Borish, Marshall Malin, Pauline Maiello, Charles A Scanga, Philana Ling Lin, Sarah M Fortune, JoAnne L Flynn

## Abstract

Immunological memory elicited either through previous or ongoing *M. tuberculosis* (Mtb) infection provides a critical mechanism by which hosts protect against re-infection and disease progression upon Mtb re-exposure. Conversely, the uneven competition between distinct Mtb strains suggest certain bacterial clades have enhanced ability to spread across communities and circulate globally, potentially by evading memory responses gained by prior infection with genomically different strains. To address whether memory responses induced by one strain can protect against a genetically distinct strain, we conducted a heterologous reinfection study in cynomolgus macaques involving primary infection by a Lineage 4 Erdman Mtb strain and subsequent re-challenge by a Lineage 2 strain, HT-L2. Recent epidemiologic studies have shown that the clade to which HT-L2 belongs has been spreading successfully over the last decade in Lima, Peru. Here, through microbiologic, PET-CT imaging and sequencing of Mtb genomic barcodes, we show that reinfected animals developed fewer lung lesions and controlled both pulmonary and disseminated forms of infection better than naïve animals without prior exposure to Mtb. Our data support that protection against reinfection is not limited by Mtb lineage, providing optimism that vaccines can be effective across populations and geographic locations.

## INTRODUCTION

Tuberculosis (TB), a primarily respiratory disease caused by infection with the slow-growing bacillus *Mycobacterium tuberculosis* (Mtb), is among the oldest infectious diseases known to afflict humans. Genetic studies have tied the pathogen to an emergence in the Horn of Africa, followed by the rise of distinct, geographically affiliated lineages that differ by hundreds to thousands of mutations.^1,2^ The geographic region to which an individual has ancestral ties has been implicated as a predictive indicator of the Mtb lineage to which an individual is most susceptible, even when infected in an area dominated by a different lineage.^3,4^ The prevalence of individual Mtb strains is certainly tied to historical events (i.e. introduction of lineages) and characteristics (e.g. density and interconnectedness with surroundings, host genetic factors) of the community, though some Mtb clades do show evidence of widespread transmission and global circulation.^5–9^ The co-evolution of the pathogen with humans has also been suggested as the source of its distinct virulence strategies,^1,10^ which enabled Mtb to survive in the human body and adapt to evade the host immune system.

Today, Mtb kills more people annually than any other infectious agent with 10.8 million estimated new cases and 1.25 million deaths in 2023.^11^ Further, a quarter of the global population is estimated to be latently or sub-clinically infected with Mtb and at potential risk of reactivation to active TB. Meta-analyses of human epidemiological data have suggested, and experimental work in animal models have corroborated, a benefit of concomitant immunity, a phenomenon in which protection is conferred by a pre-existing infection against subsequent exposures.^12,13^ Thus, investigating the breadth and mechanisms of protection against reinfection can provide critical information that is likely to be relevant for vaccines.

Recently, a genomic study of TB patients in Peru, a country conventionally dominated by Lineage 4 stains of Mtb, has identified a subclade of Lineage 2 (L2) strains that have rapidly expanded in prevalence over the last decade.^14^ Here, we refer to this subclade as high transmission (HT-L2), relative to the lower transmission strains being displaced. HT-L2 strains appear to be associated with an altered immune phenotype highlighted by lower type I interferon production, which, along with the epidemiological tracking, indicates that HT-L2 Mtb could be uniquely adapted to evade containment. Concomitant immunity and reinfection events are challenging to study in humans; degree and timing of exposure are difficult to quantify, and reinfection can only be assumed when a subject develops active TB and a strain other than the original infecting strain (if known) is detected by sputum culture. Previous studies using a model of reinfection in non-human primates (NHPs) by our group was performed with the same strain for each challenge.^13,15^ Mixed strain infections have been found in human cohorts, including combinations of Lineage 2 with non-Lineage 2 strains.^16^ Some analyses suggest that such cases, Lineage 2 strains frequently outcompete non-Lineage 2 within the same individual.^17^ Thus, we sought to determine whether its epidemiologic success indicates HT-L2 can bypass concomitant immunity and performed a heterologous strain reinfection in NHPs.

Here, Indonesian cynomolgus macaques (ICMs; *Macaca fascicularis*) were infected with a genetically barcoded library of Mtb Erdman strain (a lineage 4 strain, Erd-L4). As we could not guarantee an outcome of controlled or subclinical infection from this primary infection in all animals, this infection was cleared with anti-TB antibiotics as in our previous studies.^15,18^ This was followed by a secondary exposure to a barcoded HT-L2 strain. A parallel, Mtb-naïve group was infected with HT-L2 to compare disease outcomes without pre-existing immunity. Our results demonstrate that L4-induced immunity is capable of limiting infection and disease following subsequent exposure to HT-L2. Thus, there is cross-lineage protection induced by Mtb.

## MATERIALS AND METHODS

### Animals and husbandry

Mouse care and experimental procedures were conducted with the approval of the Institutional Animal Care and Use Committees (IACUC) of Harvard Medical School and Harvard T.H. Chan School of Public Health. The mice were housed under a 12 h light–dark cycle, ambient temperature and humidity condition in BSL3 vivarium at Harvard T.H. Chan School of Public Health.

Experiments involving macaques (*Macaca fascicularis*) were conducted at the University of Pittsburgh School of Medicine. Upon arrival, animals were quarantined, examined and confirmed to have no pre-existing Mtb infection via ELISpot assays. All experimental procedures involving care of animals complied with ethical regulations of the University of Pittsburgh School of Medicine Institutional Animal Care and Use Committee (IACUC). Macaques were pair-housed and cared for in accordance with local, state, federal, and institute policies in facilities accredited by the American Association for Accreditation of Laboratory Animal Care, under standards established in the Animal Welfare Act and the Guide for the Care and Use of Laboratory Animals as mandated by the U.S. Public Health Service Policy. Physical and behavioral health, food consumption, body weight, coughing, temperature, erythrocyte sedimentation rate, complete blood counts, and serum chemistries were monitored by veterinary staff throughout the study. Input from these examinations, along with serial PET-CT imaging, were used to determine whether criteria for the humane end point were met before the predetermined study end point.

### Barcoded HT-L2 library generation and evaluation

We selected a representative isolate from within the HT-L2 clade from a clinical strain collection that was previously defined (Peru8165).^14^ The whole genome sequence of this particular isolate can be found at BioSample: SAMN38211361 and SRA: SRS19506020. We next transformed this strain with an integrating plasmid library carrying random 18 base pair DNA sequence tags (used in Stanley et al^19^) and plated for transformants on 7H9 media with 25 μg/mL kanamycin. We then scraped up approximately 150,000 Mtb colonies to create aliquots (1 mL of OD_600_ 1.0 cells) of a master library for animal infection. To validate the complexity of the library, we extracted genomic DNA from one aliquot (1 mL of OD_600_ 1.0 cells) of the input library, PCR amplified the barcodes in the sample and performed amplicon sequencing to quantify barcode abundance on a NovaSeq10B platform, yielding approximately 1.2B reads.^20^ A modified barcode detection script to account for a 18mer barcode was run on the reads (see Github for code: https://github.com/Fortune-Lab/Mtb-WGS-barcoding) to quantify reads associated with distinct 18mer sequences.

To estimate library complexity, we ranked ordered all barcode sequences by count abundance, log-transformed the counts and then generated a linear regression model used the counts for 500 adjacent barcodes (code available on Github: https://github.com/Fortune-Lab/Mtb-WGS-barcoding). The slope from each linear model was plotted to define the difference in count abundance, with the expectation that there is a greater drop-off between counts from true barcodes and sequencing artifacts that appear due to shot noise. To define this inflection point—which represents an estimate of library complexity—we searched for a slope minima (within the first 50,000 barcodes, the maximum number of colonies we collected), which was observed at barcode rank 36,833.

### Mtb infection

Aerosol infections were performed as done previously, with some modifications.^21^ Specifically, a frozen aliquot of the HT-L2 barcoded library was thawed and diluted in PBS to 8mL of predicted OD_600_ of 0.04, which was expected to produce an infection of approximately 10 -20 bacteria per mouse. This diluted inoculum was used to aerosol infect 6-8 week old C57BL/6 female mice (The Jackson Laboratory, 000664). Mice were infected as a single batch and then randomly assigned for collection at 3 and 14 days post infection. At teach time point, mice were sacrificed and lungs homogenized for CFU enumeration and the remaining homogenate plated out as a lawn on 7H9 media supplemented with OADC for genomic DNA purification. Bacterial genomic DNA was extracted from the bacterial lawns, barcodes amplified by PCR and amplicon sequencing performed on the MiSeq (Illumina) platform.^20^ Barcode counts were quantified using the BARTI pipeline.^20^

Mtb (Erdman and HT-L2/Peru8165) was prepared similarly and instilled bronchoscopically, as previously described, targeting an inoculation dose of 15 CFU.^22,23^ The approximate inoculation dosage for each cohort was quantified by plating the bacterial suspension for CFU, with an average actual dose of ∼20 CFU (range: 6 – 33; **Supplemental Table S1**). These values are corroborated by lesion formation at four weeks post-challenge by PET CT and barcode analysis at necropsy. Challenge experiments were done at a BSL-3 facility in two cohorts with Indonesian cynomolgus macaques between 3 and 13 years of age. Cohort 1 (n = 7) was all females, while cohort 2 (n = 7) was split between males (n = 4) and females (n = 3).

### ELISpot assays

IFN-γ ELISpot assays were performed, as previously described, prior to Erd-L4 (primary) infection, between antibiotic treatment and HT-L2 (secondary) challenge, during HT-L2 infection, and at necropsy.^24^ Cryopreserved PBMCs were thawed and rested overnight in R10 media at 37°C. 2 × 10^5^ PBMCs were incubated for 40 hours with R10 media alone or a stimulation pool of ESAT-6 or CFP-10 peptides (BEI Resources) at 1 μg/mL in wells coated with IFN-γ targeted antibodies. Phorbol 12,13-dibutyrate (PDBU) and ionomycin (I; together P+I) were used as a positive control stimulation condition for each sample, guaranteeing cellular functionality after a freeze/thaw cycle. AEC peroxidase (Vector Laboratories, Inc.) colorimetrically stained spots, which were then counted on a plate reader. Samples with no conversion in the P+I condition or excessive conversion in the media alone condition were repeated with available frozen stocks, and no results are reported for samples or timepoints that were never successful. Based on historical data from our group, a result of > 10 combined spot forming units (SFU) per 2×10^5^ PBMC in response to ESAT-6 and CFP-10 was considered positive.^25^

### Macaque necropsy procedures and tissue processing

Macaques were euthanized by sodium pentobarbital injection, as previously described.^20,26^ Gross pathology observed during prosection was quantified using a published scoring system.^23,27^ PET CT scan-matched lesions were mapped and excised, along with peripheral and thoracic lymph nodes and random samples of grossly uninvolved tissue in each lung lobe. Sections of lung lodes, thoracic LNs, and large granulomas (>2 mm in diameter) were processed for formalin fixation and paraffin embedding for histological analysis. All samples were also homogenized by mechanical disruption and passing the suspension through a 70-μm cell strainer or by enzymatic dissociation (GentleMACS; Miltenyi).

### Mtb CFU determination and HT-L2 barcode analysis

Along with immunological assays, each tissue was plated for CFU quantification. Total CFU values were calculated from plating serial dilutions of homogenates on 7H11 agar plates and incubating for 21 days at 37°C in 5% CO_2_, as previously described.^23^ Granulomas detected via PET CT scan after the secondary challenge were attributed to HT-L2, as long as Erd-L4 was not found in the lesion. Granulomas (and any other CFU+ tissues) not observed on any scans were only attributed to HT-L2 if HT-L2 was detected and Erd-L4 was not.

For barcode analysis, genomic DNA was purified as done previously.^20^ Genomic DNA was then subjected to transposon-mediated fragmentation (Nextera XT, Illumina) and whole genome sequencing by the MIT BioMicroCenter (150 bp, paired end sequencing) on a NovaSeq S4 platform. All fastq reads were then passed to a custom script to detect and quantitate unique barcodes associated with both the initial Erdman Mtb library (which contained a random 7mer barcode) and the HT-L2 library, which contains 18mer barcodes. To account for less PCR bias due to limited amplification in while genome sequencing and also lower barcode sequencing depth, we removed all singleton barcodes as sequencing artifacts and considered the remainder as true barcodes in each tissue. See the following Github link for detailed description for barcode detection scripts: https://github.com/Fortune-Lab/Mtb-WGS-barcoding

### Spectral flow cytometry

Tissue homogenates were incubated for 2 hours at 37°C in 5% CO_2_ in R10 media alone or a stimulation pool of ESAT-6 and CFP-10 peptides (BEI Resources), each at 1 μg/mL. GolgiPlug (BD Biosciences) was then added at 10 μg/mL for 4 additional hours. Samples were then stained with a fixable viability dye, followed by antibody panels, detailed in **Supplemental Table S2**, for surface markers and intracellular antigens using standard protocols.^24^ Cytometry was performed at the Unified Flow Core in the University of Pittsburgh School of Medicine Department of Immunology using a five laser Aurora with SpectroFlo (16UV-16V-14B-10YG-8R; Cytek). All flow cytometry data was analyzed using FlowJo (v10.10.0) with gating strategy shown in **Supplemental Figure S2**. Unless specified otherwise, T cell populations are gated to be negative for pan-γο TCR. Due to a lack of consensus on NK cell definitions, we gated this population as CD20−CD3− lymphocytes that are CD8α+, CD16 (FcγRIIIa)+, and/or NKG2A (CD159a)+.^28^ Any samples with fewer than 100 total lymphocyte events were excluded from frequency analysis of cell types and associated functions to avoid skewing. Any samples with fewer than 50 events for a specific cell type (e.g. CD8αβ T cells) were excluded from frequency analysis of associated functions for that cell type. These samples were still included for analyses showing cell number (including cells per gram).^28^

### PET-CT scanning

All imaging occurred inside a ABSL3 housing facility according to all biosafety and radiation safety requirements at the University of Pittsburgh. Animals were weighed prior to administration of 3-5 mCi of ^18^F-fluorodeoxyglucose (FDG). Uptake of the PET tracer occurred for 50 minutes while the animals were intubated, anesthetized, and CT scanned. A mechanical ventilator held the animals’ breath for the duration (∼ 40 seconds) of the CT scan. PET-CT images were obtained with a Mediso MultiScan LFER 150 integrated preclinical PET CT and analyzed by internal image analysts using OsiriX DICOM viewer.^29–33^ Quantified features from scans include total lung FDG activity (SUVR), or the sum of all PET signal contained in the lungs above a threshold of SUV=2.3 and divided by the average PET signal in back muscle; number of granulomas, or the number of lesions observed in the lungs; and granuloma size, or the diameter of detected lesions. These measurements have been described previously.^32^

## RESULTS

### Construction and characterization of an isogenic barcoded HT-L2 library *in vitro* and in mice

To construct an isogenic wild-type barcoded library of HT-L2 for NHP infection, a representative HT-L2 clinical strain was transformed with an integrative plasmid library containing random 18mer DNA barcodes (see Methods). To benchmark this library for tracking *in vivo* infection dynamics, we assessed overall library complexity *in vitro* and infected mice to quantify early establishment infection events *in vivo* (**Figure 1A**). To ensure that each aerosol infection delivers a collection of different barcode sequences, we first assessed overall library barcode complexity by isolating Mtb genomic DNA from an aliquot of the master library, PCR amplifying the barcode sequences and subjecting the amplicons to deep sequencing. After compiling the read counts for distinct 18mer sequences, all barcodes were rank ordered by read abundance and a linear regression model used to calculate the rate of read drop-off between ranked barcodes (see Methods). We considered the greatest inflection in read abundance as the threshold between true barcodes in the library and sequencing artifacts due to variability in sampling low count barcodes or PCR chimerism. Using this approach, library complexity was calculated at approximately 36,833 distinct barcodes (**Figure 1B**). This complexity is similar to a library previously generated in Mtb Erdman, where random sampling would result in an expected 2% chance of two bacteria with the same barcode infecting a given animal in a low dose infection.^20^

**Figure 1.**
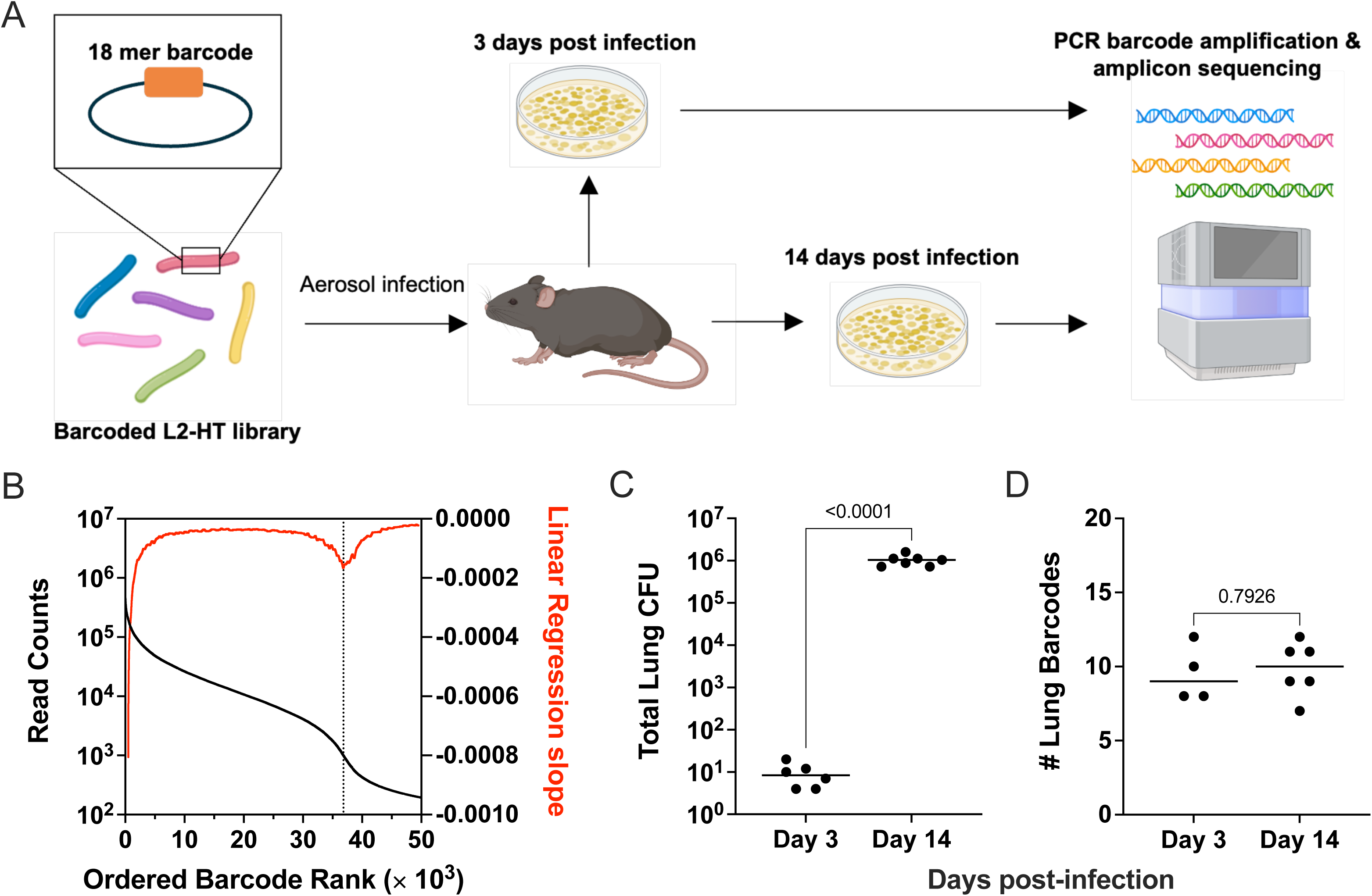
Generation and validation of genetically barcoded HT-L2 library. (**A**) Schematic of barcode design and murine validation model. C57Bl/6 mice (N = 13) were aerosol infected with an isogenic library of HT-L2 Mtb carrying random 18mer barcodes. Animals were sacrificed at 3 (n = 6) and 14 (n = 7) days post-infection and lung homogenates were plated for CFU and for Mtb genomic DNA extraction. The barcodes present in each animal were PCR amplified and quantified by amplicon sequencing. Image prepared using Biorender. (**B**) Barcodes were PCR amplified from an aliquot of the HT-L2 Mtb barcode master library and subjected to amplicon sequencing. Distinct 18mer barcodes were ordered by abundance (black curve) and library complexity was determined by performing a linear regression of log-transformed counts using a window of 500 adjacent barcodes (red curve). An inflection point, defining approximate library complexity, was defined at barcode rank 36833 (dotted line). (**C**) Total lung bacterial burden in mice infected with HT-L2 at 3- and 14-days post-challenge. (**D**) Quantification of unique barcodes recovered from lungs at 3- and 14-days post-challenge. In panels C and D, each symbol represents an animal, lines represent the group median, and p values represent the results of Welch’s unpaired t tests.

Next, we used a murine model to confirm the virulence of the barcoded HT-L2 library and validate the applicability of this library to detect early establishment events *in vivo*. C57BL/6 mice were challenged via aerosol with a low dose (target dose ∼10-20 CFU) of the HT-L2 library and lung homogenates were plated for CFU enumeration at 3 and 14 days post-infection, with lawns of bacteria from remaining homogenates also collected for Mtb genomic DNA extraction. CFU analysis showed the presence of approximately 10 bacteria in mice at 3 days post-infection, which confirmed successful low dose inoculation. We observed a significant increase in bacterial burden by 14 days post-infection, demonstrating that the barcoded library remains virulent *in vivo* (**Figure 1C**). Finally, we amplified barcodes from the genomic DNA samples and conducted amplicon sequencing to identify the number of unique barcodes present in each animal (**Figure 1D**). Approximately 10 distinct barcodes were identified in animals at both day 3 and day 14 time points, aligning closely with CFU burden at day 3. This supports that, despite a bacterial outgrowth of 5 orders of magnitude over two weeks, barcode sequencing of this library can successfully infer early Mtb establishment events *in vivo*.

### Macaque model of reinfection and establishment of primary Mtb infection

To establish a similar profile to that naturally occurring in Peru, we evaluated the efficacy of concomitant immunity conferred by a Lineage 4 infection against a subsequent challenge with the HT-L2 strain (**Figure 2A**).^14,34^ Fourteen Indonesian cynomolgus macaques (ICM) were randomly assigned to reinfection (Erd-L4 / HT-L2; n = 8) or naïve (HT-L2 only; n = 6) arms. The reinfection group was bronchoscopically infected with 20 CFU of Erd-L4, genetically barcoded to provide molecular evidence of infection that could be attributed to the primary infection when analyzed at necropsy.^13,20^ Detailed information, including infection doses for each animal, are reported in **Supplemental Table S1**.

**Figure 2.**
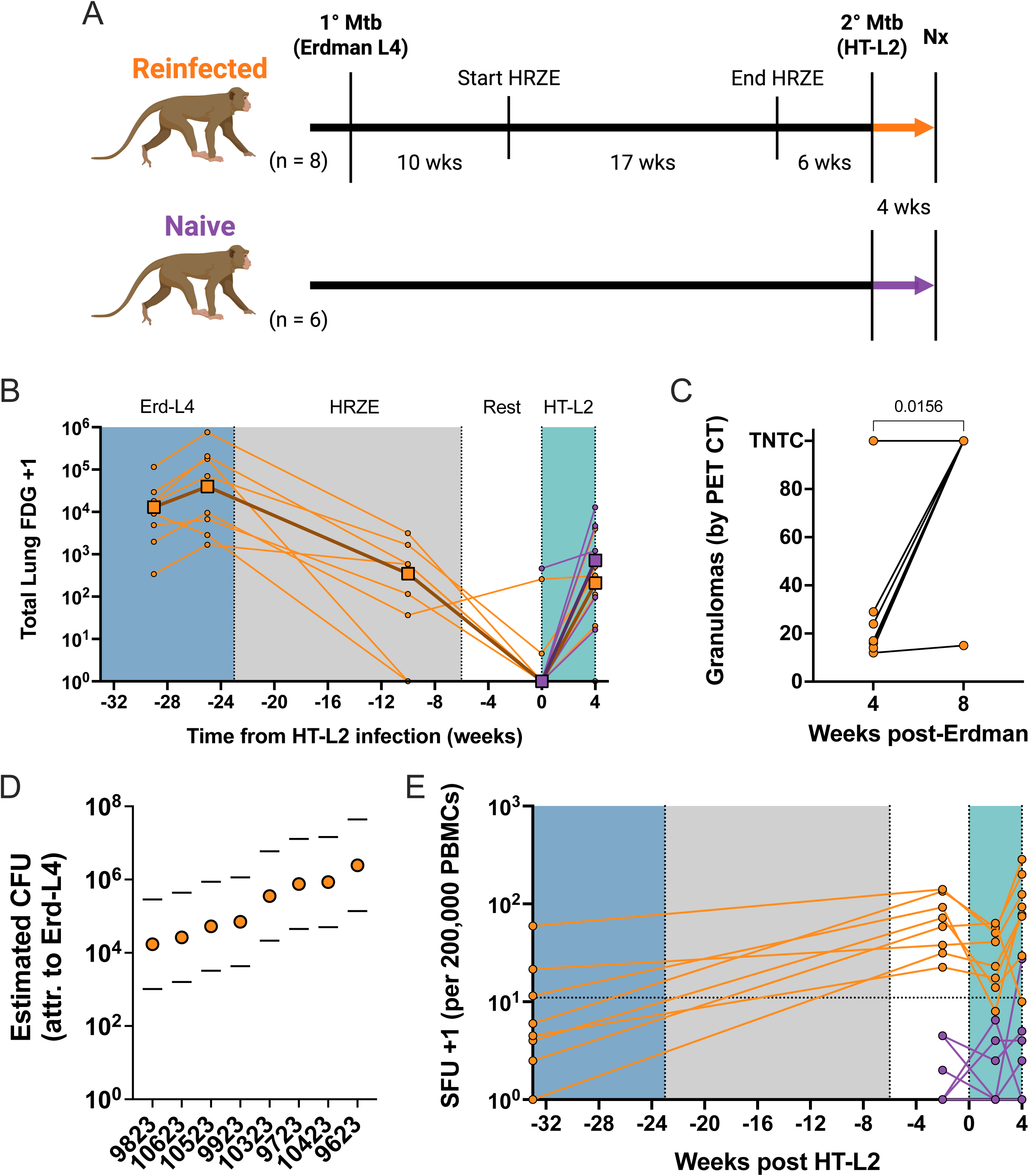
Erd-L4 infection in macaques was successfully cleared by antibiotics, although systemic memory responses remained intact. (**A**) Schematic of macaque reinfection timeline. (**B**) Quantification of inflammation in lung via serial PET scans throughout primary infection (Erd-L4, blue shading), antibiotic treatment (HRZE, gray shading), and secondary infection (HT-L2, green shading). Thin lines with circles represent individual animals (orange Erd-L4 then HT-L2; purple HT-L2 in naïve animals); thick lines with squares represent group median. (**C**) Number of granulomas appearing on PET CT scans at 4 and 8 weeks post-primary Erd-L4 infection. Each symbol represents an animal and the p value is the result of a Wilcoxon signed rank test. (**D**) Estimated thoracic bacterial burden per animal at 8 weeks post-primary Erd-L4 infection based on PET CT scans, as described in Refs. ^13^ and ^27^. Circles represent point estimate; Lines represent extent of 95% confidence interval. (**E**) IFN-g release in response to stimulation with ESAT-6 and CFP-10 peptide pools was quantified by ELISpot assay throughout reinfection study. A threshold of 10 spot forming units (SFU) per 2 × 10^5^ PBMCs for positive results was determined by previous studies (Ref. ^25^). Lines connect timepoints for each individual, and each symbol represents and animal.

We employed serial positron emission tomography and computed tomography (PET CT) scans in parallel with clinical observations to confirm establishment and progression of the primary Erd-L4 infection.^32^ Scan analysis confirmed establishment of active infection by four weeks post-challenge (**Figure 2B**) and primary infection lesions were annotated as they formed.^20,22,32^ Disease progressed significantly by 8 weeks post-infection, with 88% (7/8) of animals developing large consolidations across multiple lung lobes and estimated (based on total lung FDG activity) thoracic bacterial burdens as high as 2.5 × 10^6^ (**Figure 2C,D**) indicating poor initial control of the Erd-L4 strain in the ICMs.^13,27^

As in previous reinfection studies, the primary Erd-L4 infection was then treated beginning at 10 weeks post-infection with a standard four-month regimen of anti-tuberculous drugs (rifampicin, isoniazid, pyrazinamide, ethambutol; HRZE) at human-equivalent doses.^15,18^ PET CT scans confirmed that lung inflammation returned to low levels that were comparable to the naïve control animals following the HRZE regimen (**Figure 2B**).

We confirmed the development of an adaptive anti-Mtb response in Erd-L4 infected ICMs via an IFN-γ enzyme linked immunosorbent spot (ELISpot) assay to quantify the frequency of T cells in blood releasing IFNγ following stimulation with Mtb antigens ESAT6 and CFP10. All 8 Erd-L4 challenged macaques converted to positive (median: 64.25 SFU) prior to HT-L2 challenge, indicating that a systemic response was successfully mounted prior to, and persisted beyond, antibiotic therapy (**Figure 2E**). Naïve ICMs remained negative (median: 0.5 SFU) at the pre-HT-L2 timepoint.

### HT-L2 challenge successfully initiated active TB disease in naïve macaques

At 33 weeks post-Erd-L4 (6 weeks after cessation of HRZE), the 8 Erd-L4 macaques and the 6 naïve macaques were bronchoscopically challenged with the barcoded strain of HT-L2 (median dose: 19.5 CFU) and necropsied one month later (median time to necropsy: 5.1 weeks). PET CT imaging performed prior to necropsy assessed the extent of new disease following HT-L2 challenge. Total lung FDG activity was similar between reinfected and naïve groups at this timepoint (**Figure 2B**). ELISpot assays 2 weeks post-secondary infection and at time of necropsy showed a lack of Mtb-specific response mounted in the naïve group, which aligns with delayed mounting of adaptive responses and conversion timeframe in early active TB compared to other infections (**Figure 2E**).^24,35,36^

### Previous Mtb infection provides cross-lineage immunity

Comprehensive necropsies were performed ∼4 weeks post HT-L2 challenge. The pre-necropsy PET CT scan was used as a map to identify and isolate individual granulomas and other pathologic lesions. Using the full set of scans across both infections, lesions were identified as “old” from the Erd-L4 infection or “new” if appearing post-HT-L2 infection. Each scan-matched lesion was excised along with multiple random sections of each lung lobe and all peripheral and thoracic lymph nodes. There were significantly fewer “new” granulomas that were attributable to HT-L2 infection (determined via PET CT and verified by sequencing for barcode analyses) within the reinfected group compared to the naïve group (**Figure 3A**). Barcode quantification of each CFU+ sample (granulomas, lymph nodes, lungs) determined that significantly fewer HT-L2 barcodes were found in animals that had been previously challenged with Erd-L4 (**Figure 3B**). Together, these data suggest that immune pressure induced by the previous infection serves as a restrictive bottleneck that impedes establishment of the second infection.

**Figure 3.**
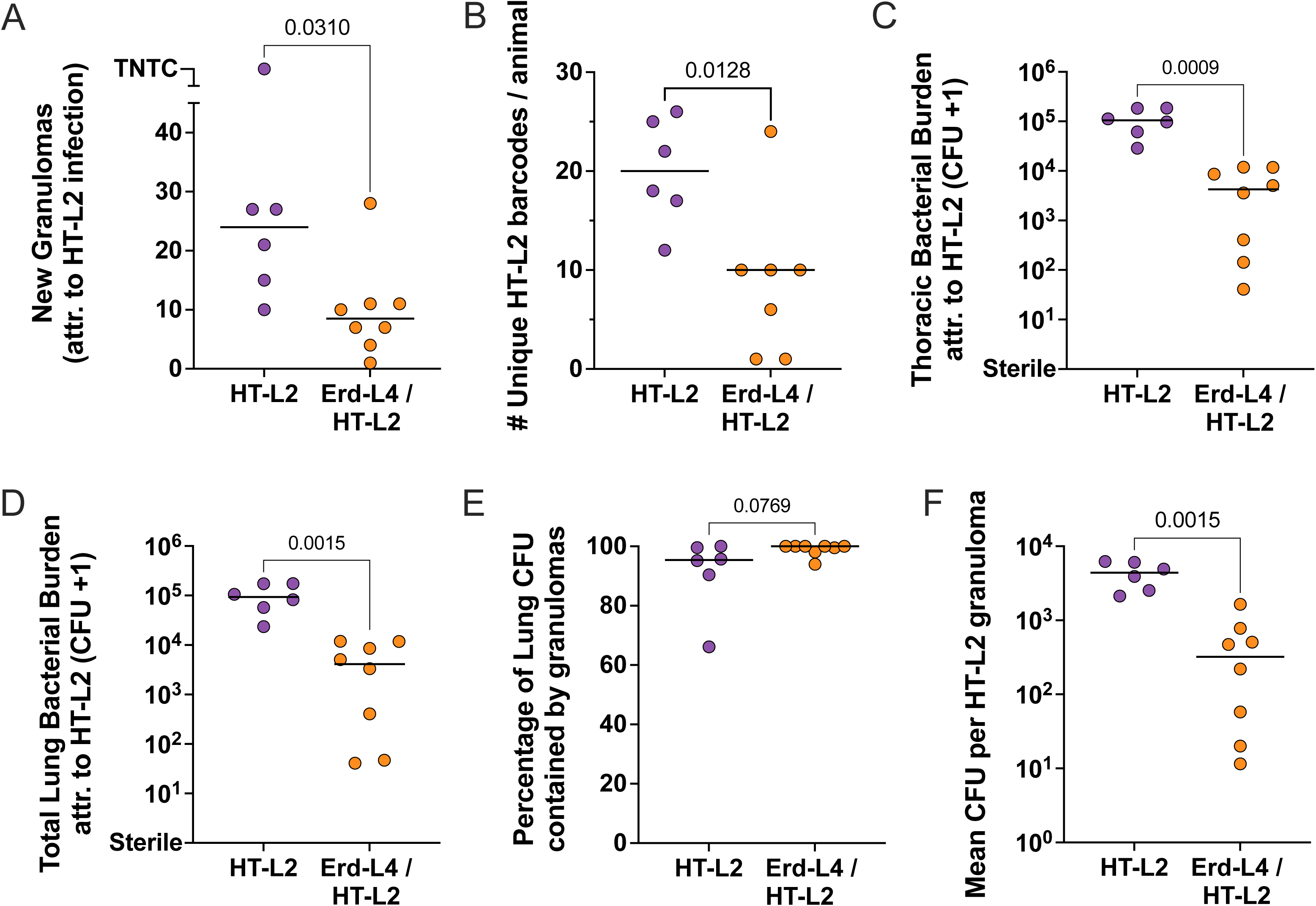
Animals previously infected with Erd-L4 were significantly less susceptible to HT-L2 infection. (**A**) Number of granulomas found at necropsy that were clinically and microbiologically attributable to HT-L2 infection. (**B**) Number of unique HT-L2 barcodes recovered from each animal. One animal (10623) is excluded as no sequencing samples passed QC. (**C,D**) Bacterial burden attributable to HT-L2 in all thoracic (C) and only lung (D) tissues. (**E**) Percentage of lung bacterial burden obtained from isolated granulomas, rather than in random samples of grossly uninvolved lung tissue, as an indicator of successful granuloma containment. This was calculated by dividing the summed granuloma-attributed CFU by the lung CFU in panel D. (**F**) Average HT-L2 bacterial burden per granuloma in reinfected and naïve animals. In all plots, each symbol represents an animal and lines represent the group median. Statistics: P-values shown with Mann-Whitney non-parametric ranked sum tests in panels A, B, and E, and Welch’s unpaired t tests for panels C, D and F.

The reinfected group also had significantly reduced total thoracic bacterial burden (∼50-fold) attributable to the HT-L2 infection compared to the naïve group, which was the predetermined primary outcome measure in the study (**Figure 3C**). Erd-L4 was recovered from 2 thoracic lymph node sites in one animal, and these CFU were excluded from bacterial burden values. Bacterial burden in the lung, consisting of CFU from granulomas and random samples of grossly uninvolved lung lobe, was significantly reduced compared to the naïve controls (**Figure 3D**). A higher percentage of the lung CFU was recovered from granulomas (rather than non-grossly involved lung) from reinfected compared to naïve animals, which suggests less granuloma failure, reduced intra-lung dissemination and formation of fewer early-stage lesions that were not detectable macroscopically or on scans (**Figure 3E**). The CFU+ granulomas stemming from the secondary infection in reinfected macaques had a lower average bacterial burden, providing further indication of improved containment or killing of those HT-L2 Mtb that did establish infection, presumably due to protective immune responses induced by the Erd-L4 infection (**Figure 3F**). In one animal, HT-L2 was recovered from an “old” lesion that formed following the primary infection with Erd-L4, confirmed by PET CT, and was subsequently cleared of Erd-L4 by antibiotic treatment. This phenomenon of trafficking infecting bacteria to previous sites of infection has been observed previously in this model.^13,15^

### Mtb strains and exposure history lead to variability in lesion characteristics

We used PET CT scans to evaluate the size of granulomas at 4 weeks post-infection in macaques exposed to both Erd-L4 and HT-L2. Granulomas at 4 weeks attributable to HT-L2 were appreciably smaller than those from Erd-L4 at 4 weeks in most animals (**Supplemental Figure S1A**). However, comparison of HT-L2 lesions in the naïve cohort with those in the reinfected animals showed negligible size differences (**Supplemental Figure S1B**). Since granulomas attributable to the secondary infection had significantly lower bacterial burden than those in primary disease (**Figure 3F**), granuloma size does not appear to primarily indicate protective capacity of the lesion (**Supplemental Figure S1B**) and may be influenced by the causative strain (**Supplemental Figure S1A**). Granulomas at time of necropsy were also evaluated for 18F-FDG avidity via PET, a marker of inflammation; 38% of the reinfection animals (3/8) had no FDG avid lesions compared to 0/6 of naïve animals (**Supplemental Figure S1C**).

We employed spectral flow cytometry to evaluate the lymphocyte composition within individual HT-L2 granulomas. This analysis revealed that the relative abundance of various cell types remained largely consistent among lesions, animals and groups (**Figure 4A**). Granulomas in reinfected animals had fewer cells per lesion on average than those in naïve animals (**Figure 4B**). This reduction in cell numbers was primarily driven by a significant reduction in γδ T cells (p = 0.0252) and reductions in NK (p = 0.0958) and conventional T cell (p = 0.0565) populations (**Figure 4C**). Of the conventional T cell subsets, classical CD8αβ T cells (p = 0.0523) showed the strongest trend towards a decrease, although CD4 (p = 0.1135) and CD8αα (p = 0.1307) T cells appear to be following a similar pattern (**Figure 4D**). This may represent that granulomas in reinfection have successfully limited bacterial replication and begun transitioning from a pro-inflammatory to a wound healing environment, as seen in our previous reinfection study.^15^

**Figure 4.**
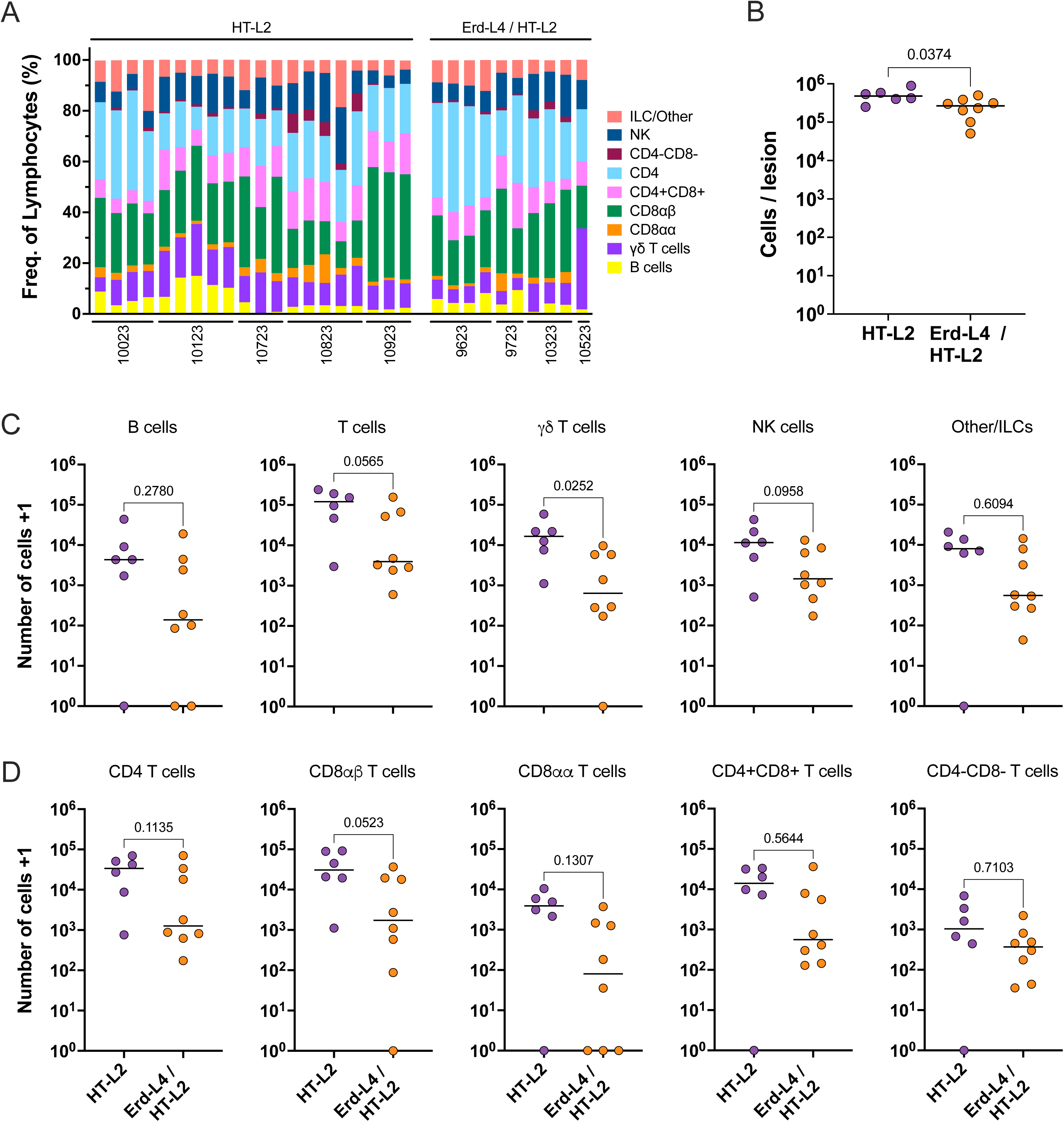
Granulomas of reinfected animals had fewer cells, driven by a drop in traditionally cytotoxic cell types. (**A**) Relative frequency of lymphocyte subsets by flow cytometry. Each bar represents a granuloma analyzed from the animal noted below. (**B**) Average number of total cells recovered per lesion by animal, as determined by hemocytometer. Lesions with cell counts below limit of detection on hemocytometer (2 × 10^4^ cells) were excluded. (**C,D**) Average number of cells, broken down by lymphocyte (C) and T cell (D) subsets, in granulomas analyzed by flow cytometry. In panels B-D, each symbol represents an animal, lines represent the group median, and all p values shown are results of Welch’s unpaired t tests.

### Protection from reinfection is associated with an altered immune profile

The immune profile of lung tissue is modified by previous Mtb infection. The frequency of lung CD8αβ T cell populations producing cytotoxic effector molecules significantly increased, with a trend (p = 0.0813) also seen in increasing cytotoxic CD4 T cells (**Figure 5A**). Both conventional T cell subsets in granulomas significantly shifted towards increased production of cytokines traditionally associated with anti-mycobacterial immune responses (**Figure 5B**). Of note, there was a trend (p = 0.0695) towards more granzyme K+ CD4 T cells/gram lung tissue in macaques with prior infection compared to naïve macaques following HT-L2 infection (**Figure 5C**). Granzyme K+ CD8 T cells identified by single cell RNA sequencing analysis of granulomas in a previous reinfection study were associated with protection against disease from the secondary exposure; this population was greatly reduced by CD4 depletion.^15^ Other pro-inflammatory responses, like granulysin and IFN-γ production, trended higher in the reinfection group. No apparent differences in the number of effector populations were observed in the CD8 T cells (**Figure 5D**). This demonstrates a balance between the efficiency of memory responses and amplitude of ineffective naïve responses (**Figures 4B, 5A-D**). Overall, these results suggest that the anti-mycobacterial responses are more robust and efficient in macaques with prior infection, likely resulting in increased containment of Mtb and protection against disease.

**Figure 5.**
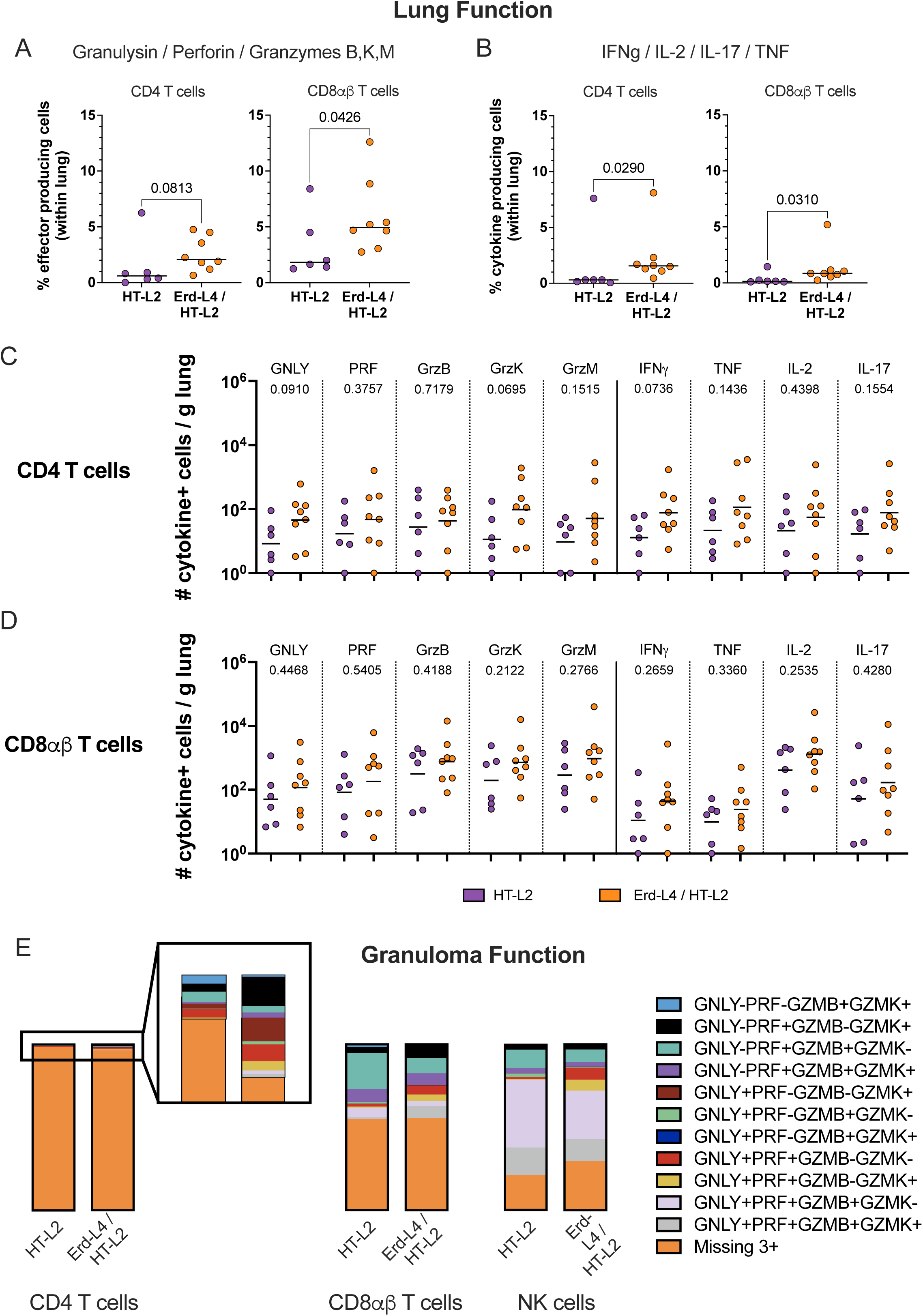
T cell populations were more efficient at producing functional responses, with a shift towards granzyme K positive phenotypes. (**A,B**) Frequency of CD4 and CD8αβ T cells in lung tissue producing cytotoxic effectors (A) and cytokines (B). p values reported for Mann-Whitney non-parametric ranked sum tests between groups. (**C,D**) Number of CD4 (C) and CD8αβ (D) T cells per gram of lung tissue producing individual effector molecules. p values reported for Welch’s unpaired t tests, with groups compared for each effector. In panels A-D, each symbol represents an animal and lines represent the group median. (**E**) Frequency of CD4 T, CD8αβ T and NK cells in lung granulomas producing competent cytotoxic responses, determined by production of at least two of granulysin, perforin, granzyme B and granzyme K.

Cytotoxic effector function is complex with multiple simultaneous functions (e.g. co-production of granzymes with perforin or granulysin) required for efficacy.^37–39^ Within the reinfection granuloma specifically, effector CD8αβ and CD4 T cell populations appeared to shift towards granzyme K producing phenotypes, as seen by expansion of the GNLY-PRF+GZMB-GZMK+, GNLY+PRF+GZMB-GZMK+ and GNLY+PRF+GZMB+GZMK+ populations (black, yellow and grey bands, respectively) and shrinking of GNLY-PRF+GZMB-GZMK+ and GNLY+PRF+GZMB+GZMK-populations (teal and pink bands, respectively) (**Figure 5E**). In both CD8αβ T cells and NK cells, a relative increase in granulysin/perforin double positive cells that do not produce either granzymes B or K (red band) was observed among reinfected animals.

### Dissemination of secondary infections is well contained

Another clinically relevant feature associated with protection is prevention of dissemination. Initial dissemination to lymph nodes is a normal event following Mtb infection, which results in priming of Mtb specific T cell responses.^40–43^ However, increased dissemination within the lungs and to thoracic lymph nodes is a sign of progressive disease.

We combined Mtb strain barcode analysis with PET CT imaging (**Figure 3A**) and quantitative microbiology (**Figure 3C-F**) to assess dissemination of the HT-L2 strain in macaques. Prior infection with Erd-L4 provided significant protection against dissemination of the secondary HT-L2 infection, as barcodes from HT-L2 strain were shared across fewer tissues in reinfected macaques compared to naïve macaques (**Figure 6A**). In the macaque model, TB dissemination is most common within lungs and to lung draining lymph nodes.^43^ There was significantly lower bacterial burden in thoracic lymph nodes of reinfected animals, with half of the animals maintaining sterile thoracic lymph nodes (**Figure 6B,C**). One macaque (10323) had Erd-L4 barcodes recovered from two thoracic lymph nodes, while another (10223) had both Erd-L4 and HT-L2 barcodes recovered from a peripheral lymph node. This is not surprising since lymph nodes are known reservoirs of persistent Mtb even following drug treatment.^15,18,42,43^ At the individual lymph node level, there was considerable Mtb growth in lymph nodes in naive animals, while the relatively few CFU lymph nodes in reinfected animals maintained very low bacterial burdens. (**Figure 6D**). There was significantly less lung-to-lymph node dissemination events in the reinfection animals compared to naïve animals (**Figure 6E**). This could be a direct result of fewer lung granulomas establishing and improved control in those granulomas and/or better control in distal sites (i.e. lymph nodes) that prevent outgrowth (and barcode detection). Together these data demonstrate a significant restriction of HT-L2 dissemination to and growth in lymph nodes, suggesting that immune containment likely occurs both at the original infection sites (i.e. airways and lung), thereby preventing dissemination, and within the lymph node by limiting growth or increasing killing.

**Figure 6.**
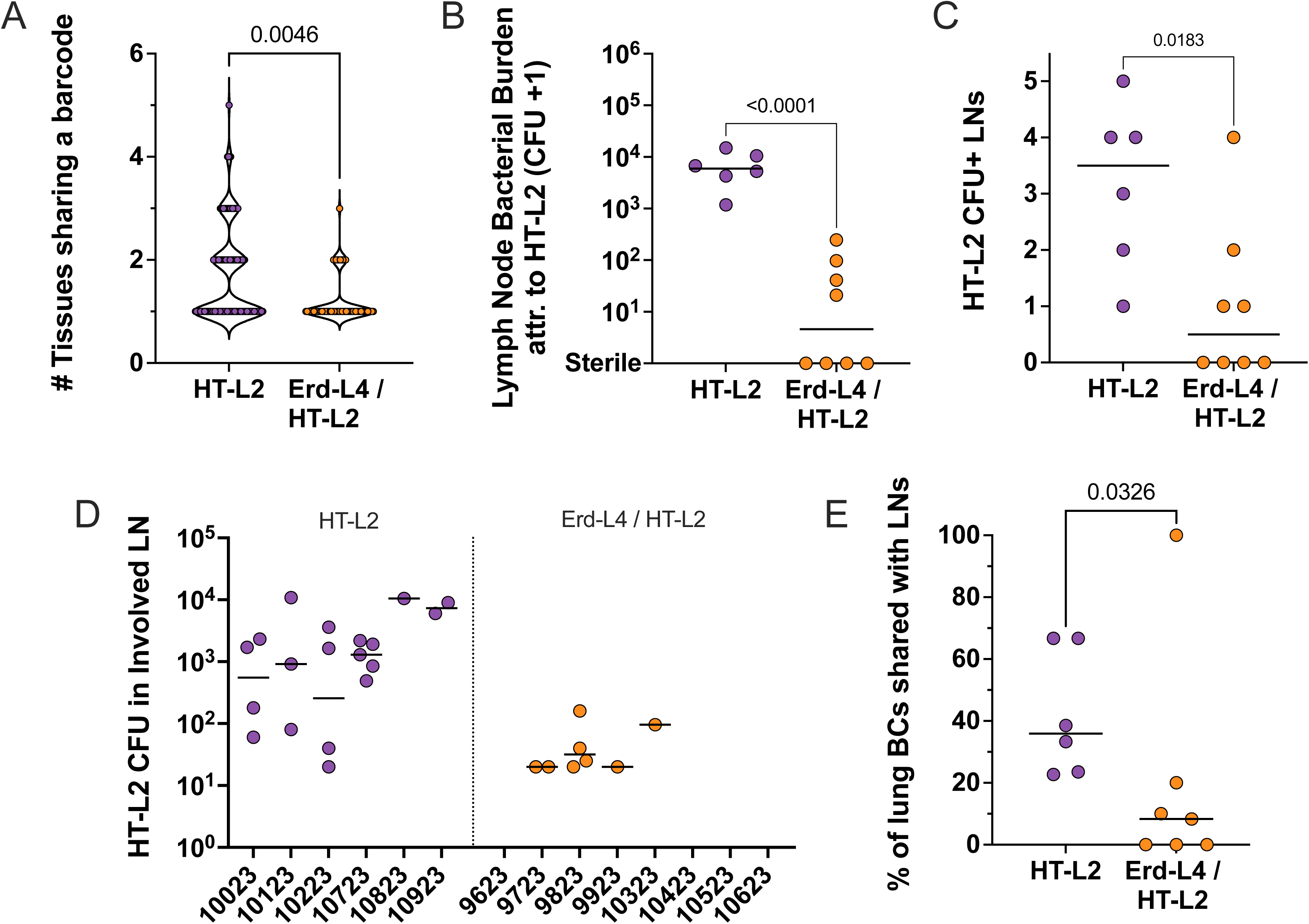
Pre-existing immunity from a lineage 4 infection limits dissemination of subsequent lineage 2 infections. (**A**) Extent of overall dissemination, shown by the number of tissues sharing a barcode. Each symbol represents a barcode. Reported p value shown for Kolmogorov-Smirnov test. (**B**) Bacterial burden attributable to HT-L2 infection in lymph node tissues. Reported p value shown for Welch’s unpaired t test. (**C**) The number of CFU positive lymph nodes per animal. (**D**) Bacterial burden per CFU+ lymph node, shown by animal. Each symbol represents a lymph node, and lines represent the median per animal. (**E**) Percentage of barcodes recovered from a lung tissue that are shared with a lymph node site of dissemination. In panels B, C and E, each symbol represents an animal and lines represent the group median. In panels C and E, p values are results of Mann-Whitney non-parametric ranked sum test. In panels A and E, one animal (10623) is excluded as no sequencing samples passed QC.

## DISCUSSION

Most non-human primate models of TB employ the laboratory strain Mtb Erdman, although others, such as H37Rv, are occasionally used.^44–46^ This has been beneficial to the field as results across studies can be compared more easily. However, strain and lineage differences are well documented in humans. Thus, it is important that pre-clinical research reflects clinical and immunological diversity by implementing a wider variety of Mtb strains. While clinical isolates (e.g. HN878, CDC1551 and SA161) have been used to various extents in murine models, their adoption for NHP work has been more limited and, in the case of CDC1551, there has been controversy over virulence qualities in NHP.^47–50^ Here, the barcoded Peruvian HT-L2 strain was shown to successfully establish infection in both mice and macaques, and will serve as a useful tool for studying clinical isolates.

Concomitant immunity is a fundamental aspect of TB that remains poorly understood despite the potential for influencing vaccine strategies, clinical trial design, and epidemiology.

Understanding the underlying immune features, mechanisms, and limitations of protection conferred by prior exposure to Mtb is crucial to furthering our development of prophylactic and therapeutic interventions and, ultimately, minimizing global TB burden. Here, we demonstrated that the protective immunity from prior Mtb infection is robust and reduces disease burden resulting from exposures to a genetically distinct lineage and strain known to illicit a unique immune profile.^14^ Protection was observed by several metrics: total thoracic, lung and lymph node bacterial burden due to HT-L2 were all significantly lower in reinfected macaques compared to naïve macaques, with reduced establishment and dissemination of HT-L2 in reinfected compared to naïve macaques. Of note, the magnitude of protection was similar to our previous homologous strain (Erd-L4/Erd-L4) reinfection studies where initial infection was treated with anti-TB drugs.^15,18^ As seen previously, drug treatment reduces, but does not abolish, protection against reinfection.^18^

The primary interpretation of these data is that there are common immunogenic antigens conserved across Mtb strains and lineages which contribute to protective memory responses. The adaptive arm of the immune system is a significant contributor to concomitant immunity, and this protection is, at least in part, CD4 T cell dependent.^15^ Overall, this conclusion indicates that a universal vaccine – that is, efficacious irrespective of patient ancestry, case location or infection strain – against Mtb is possible. However, these results also suggest that we have yet to identify which, or perhaps how many, antigens are critical for triggering protective immunity. A vaccine strategy that induces durable breadth of protective responses similar to prior infection may be required. It is also possible that primary memory responses in effective models of disease prevention, such as IV BCG vaccination, CMV expressing Mtb antigens and Mtb reinfection, are different and rely on distinct sets of antigens.^13,24,45^ Nonetheless, prolonged exposure to a breadth of antigens and, apparently, durable immune responses that are protective are common features among these vaccine strategies.

Our immunology data on heterologous reinfection aligns with previous data in reinfection models supporting the notion that a balance of pro-inflammatory and equally important anti-inflammatory responses are key to rapid control of infection. The role of granzyme K secreting T cells earlier than four weeks post-infection should be investigated to determine whether they are important in initial control or are part of the healing response. The immunology data, in combination with barcoding analysis confirming reduced establishment and early bacterial restriction following reinfection, suggest that the protective responses can rapidly contain the infection and begin resolving lesions within four weeks.

It is not clear at this point how generalizable our findings are. Concurrent infection with multiple strains has been observed in humans, but several questions remain unanswered regarding the phenomenon of concomitant immunity in the context of heterologous re-challenge. Does the order of strain infections influence protection? That is, would HT-L2 primary infection lead to protective responses against Erd-L4, or even HT-L2 itself? Would a lower transmission strain, such as the Lineage 2 strains being outcompeted in Peru, provide little protection and/or be easily protected against? It is possible that Erd-L4 primary infection leads to broadly protective immunity, although a single experiment cannot discount the potential for strains that are capable of escaping containment from Erd-L4 induced concomitant immunity. If universal protection is obtainable, the mechanisms (i.e. innate training vs adaptive memory) or contributing antigens (i.e. highly conserved mycolipids, implicating innate-like subsets, or peptides, implicating conventional T cells) must be elucidated to benefit the development pipelines of interventions. However, a scenario in which concomitant immunity against heterologous reinfection is not universally protective may also serve as an opportunity to determine why specific combinations, such as Erd-L4 / HT-L2, lead to protection while other do not by comparing antigenicity and expanded TCR or BCR profiles. NHPs can provide a model system for testing specific strain pairings that may be relevant to vaccine or therapeutic trial design.

Limitations of this study include the use of Indonesian cynomolgus macaques, due to the lack of availability of Chinese cynomolgus macaques, thereby limiting the ability to directly compare against our historical data on reinfection models. Larger samples sizes of macaques would likely provide clearer immunologic differences, although the sample size calculated *a priori* was sufficient to obtain significant results in our primary and secondary outcomes (CFU and barcode establishment). Since fewer granulomas establish in cases of reinfection, the sample size for comparison with naïve animals is limited. This was also the first study to infect NHPs with the HT-L2 strain isolated in Peru and we have no data on HT-L2 outcomes over a longer infection duration. Additional studies on the infection trajectory and protection against reinfection with a wider range of clinical Mtb strains will be necessary to fully understand the effects of Mtb strain on immunity.

## ACKNOWLEDGEMENTS

This work was supported by the National Institutes of Health [IMPAc-TB (HI-IMPACT) 75N93019C00071, T32 5T32AI089443, T32 5T32AI060525]; and the Bill and Melinda Gates Foundation, Seattle, WA [Gates Foundation Investment Award INV-020435].

## DATA AVAILABILITY

All amplicon and whole genome sequencing data in this work is publicly available on SRA under the accession number: PRJNA1290335.

## REFERENCES

1 Comas, I. et al. Out-of-Africa migration and Neolithic coexpansion of Mycobacterium tuberculosis with modern humans. Nature Genetics 45, 1176–1182 (2013). 10.1038/ng.2744

2 Bañuls, A. L., Sanou, A., Van Anh, N. T. & Godreuil, S. Mycobacterium tuberculosis: ecology and evolution of a human bacterium. J Med Microbiol 64, 1261–1269 (2015). 10.1099/jmm.0.000171

3 Hirsh, A. E., Tsolaki, A. G., Deriemer, K., Feldman, M. W. & Small, P. M. Stable association between strains of *Mycobacterium tuberculosis* and their human host populations. Proceedings of the National Academy of Sciences 101, 4871–4876 (2004). 10.1073/pnas.0305627101

4 Gagneux, S. et al. Variable host–pathogen compatibility in *Mycobacterium tuberculosis*. Proceedings of the National Academy of Sciences 103, 2869–2873 (2006). 10.1073/pnas.0511240103

5 Dou, H. Y., Huang, S. C. & Su, I. J. Prevalence of Mycobacterium tuberculosis in Taiwan: A Model for Strain Evolution Linked to Population Migration. Int J Evol Biol 2011, 937434 (2011). 10.4061/2011/937434

6 Hsu, Y. H. et al. Association of NRAMP 1 gene polymorphism with susceptibility to tuberculosis in Taiwanese aboriginals. J Formos Med Assoc 105, 363–369 (2006). 10.1016/S0929-6646(09)60131-5

7 Aravindan, P. P. Host genetics and tuberculosis: Theory of genetic polymorphism and tuberculosis. Lung India 36, 244–252 (2019). 10.4103/lungindia.lungindia_146_15

8 Bellamy, R. Genetics and pulmonary medicine bullet 3: Genetic susceptibility to tuberculosis in human populations. Thorax 53, 588–593 (1998). 10.1136/thx.53.7.588

9 Gagneux, S. Host–pathogen coevolution in human tuberculosis. Philosophical Transactions of the Royal Society B: Biological Sciences 367, 850–859 (2012). 10.1098/rstb.2011.0316

10 Wolfe, N. D., Dunavan, C. P. & Diamond, J. Origins of major human infectious diseases. Nature 447, 279–283 (2007). 10.1038/nature05775

11 Global tuberculosis report. (World Health Organization, Geneva, 2024).

12 Andrews, J. R. et al. Risk of Progression to Active Tuberculosis Following Reinfection With Mycobacterium tuberculosis. Clinical Infectious Diseases 54, 784–791 (2012). 10.1093/cid/cir951

13 Cadena, A. M. et al. Concurrent infection with Mycobacterium tuberculosis confers robust protection against secondary infection in macaques. (2021). 10.1371/journal.ppat.1007305

14 Luo, Y. et al. Paired analysis of host and pathogen genomes identifies determinants of human tuberculosis. Nature Communications 15 (2024). 10.1038/s41467-024-54741-w

15 Bromley, J. D. et al. CD4+ T cells re-wire granuloma cellularity and regulatory networks to promote immunomodulation following Mtb reinfection. Immunity 57, 2380–2398.e2386 (2024). 10.1016/j.immuni.2024.08.002

16 Warren, R. M. et al. Patients with active tuberculosis often have different strains in the same sputum specimen. Am J Respir Crit Care Med 169, 610–614 (2004). 10.1164/rccm.200305-714OC

17 Wang, J.-Y. et al. Mixed infection with Beijing and non-Beijing strains in pulmonary tuberculosis in Taiwan: prevalence, risk factors, and dominant strain. Clinical Microbiology and Infection 17, 1239–1245 (2011). 10.1111/j.1469-0691.2010.03401.x

18 Ganchua, S. K. et al. Antibiotic treatment modestly reduces protection against *Mycobacterium tuberculosis* reinfection in macaques. Infection and Immunity 92 (2024). 10.1128/iai.00535-23

19 Stanley, S. et al. Identification of bacterial determinants of tuberculosis infection and treatment outcomes: a phenogenomic analysis of clinical strains. Lancet Microbe 5, e570–e580 (2024). 10.1016/S2666-5247(24)00022-3

20 Martin, C. et al. Digitally Barcoding *Mycobacterium tuberculosis* Reveals In Vivo Infection Dynamics in the Macaque Model of Tuberculosis. mBio 8, e00312–00317 (2017). 10.1128/mBio.00312-17

21 Wang, X. et al. Engineered Mycobacterium tuberculosis triple-kill-switch strain provides controlled tuberculosis infection in animal models. Nature Microbiology 10, 482–494 (2025). 10.1038/s41564-024-01913-5

22 Capuano, S. V. et al. Experimental *Mycobacterium tuberculosis* Infection of Cynomolgus Macaques Closely Resembles the Various Manifestations of Human *M. tuberculosis* Infection. Infection and Immunity 71, 5831–5844 (2003). 10.1128/iai.71.10.5831-5844.2003

23 Lin, P. L. et al. Quantitative Comparison of Active and Latent Tuberculosis in the Cynomolgus Macaque Model. Infection and Immunity 77, 4631–4642 (2009). 10.1128/iai.00592-09

24 Darrah, P. et al. Prevention of tuberculosis in macaques after intravenous BCG immunization. Nature 577, 95–102 (2020). 10.1038/s41586-019-1817-8

25 Darrah, P. A. et al. Airway T cells are a correlate of i.v. Bacille Calmette-Guerin-mediated protection against tuberculosis in rhesus macaques. Cell Host & Microbe 31, 962–977.e968 (2023). 10.1016/j.chom.2023.05.006

26 Lin, P. L. et al. Sterilization of granulomas is common in active and latent tuberculosis despite within-host variability in bacterial killing. Nature Medicine 20, 75–79 (2013). doi:10.1038/nm.3412

27 Maiello, P. et al. Rhesus Macaques Are More Susceptible to Progressive Tuberculosis than Cynomolgus Macaques: a Quantitative Comparison. Infection and Immunity 86, e00505–00517 (2021). 10.1128/IAI.00505-17

28 Simonson, A. W. et al. Intravenous BCG-mediated protection against tuberculosis requires CD4+ T cells and CD8α+ lymphocytes. J Exp Med 222 (2025). 10.1084/jem.20241571

29 Sarnyai, Z. et al. Performance Evaluation of a High-Resolution Nonhuman Primate PET/CT System. Journal of Nuclear Medicine 60, 1818–1824 (2019). 10.2967/jnumed.117.206243

30 Rosset, A., Spadola, L. & Ratib, O. OsiriX: An Open-Source Software for Navigating in Multidimensional DICOM Images. Journal of Digital Imaging 17, 205–216 (2004). 10.1007/s10278-004-1014-6

31 Coleman, M. T. et al. Early Changes by 18Fluorodeoxyglucose Positron Emission Tomography Coregistered with Computed Tomography Predict Outcome after Mycobacterium tuberculosis Infection in Cynomolgus Macaques. Infection and Immunity 82, 2400–2404 (2014). 10.1128/iai.01599-13

32 White, A. G. et al. Analysis of 18FDG PET/CT Imaging as a Tool for Studying *Mycobacterium tuberculosis* Infection and Treatment in Non-human Primates. JoVE Immunologu and Infection, 56375 (2017). 10.3791/56375

33 Diedrich, C. R. et al. SIV and Mycobacterium tuberculosis synergy within the granuloma accelerates the reactivation pattern of latent tuberculosis. PLOS Pathogens 16, e1008413 (2020). 10.1371/journal.ppat.1008413

34 Barletta, F. et al. Predominant Mycobacterium tuberculosis Families and High Rates of Recent Transmission among New Cases Are Not Associated with Primary Multidrug Resistance in Lima, Peru. Journal of Clinical Microbiology 53, 1854–1863 (2015). 10.1128/jcm.03585-14

35 Menzies, D. Interpretation of repeated tuberculin tests. Boosting, conversion, and reversion. Am J Respir Crit Care Med 159, 15–21 (1999). 10.1164/ajrccm.159.1.9801120

36 Poulsen, A. Some clinical features of tuberculosis. 1. Incubation period. Acta Tuberc Scand 24, 311–346 (1950).

37 Andersson, J., Samarina, A., Fink, J., Rahman, S. & GrundströM, S. Impaired Expression of Perforin and Granulysin in CD8+ T Cells at the Site of Infection in Human Chronic Pulmonary Tuberculosis. Infection and Immunity 75, 5210–5222 (2007). 10.1128/iai.00624-07

38 Voskoboinik, I., Whisstock, J. C. & Trapani, J. A. Perforin and granzymes: function, dysfunction and human pathology. Nature Reviews Immunology 15, 388–400 (2015). 10.1038/nri3839

39 Pitabut, N., Dhepakson, P., Sakurada, S., Keicho, N. & Khusmith, S. Coordinated In Vitro Release of Granulysin, Perforin and IFN-γ in TB and HIV/TB Co-Infection Associated with Clinical Outcomes before and after Anti-TB Treatment. Pathogens 9, 655 (2020). 10.3390/pathogens9080655

40 Chackerian, A. A., Alt, J. M., Perera, T. V., Dascher, C. C. & Behar, S. M. Dissemination of *Mycobacterium tuberculosis* Is Influenced by Host Factors and Precedes the Initiation of T-Cell Immunity. Infection and Immunity 70, 4501–4509 (2002). 10.1128/iai.70.8.4501-4509.2002

41 Wolf, A. J. et al. Initiation of the adaptive immune response to *Mycobacterium tuberculosis* depends on antigen production in the local lymph node, not the lungs. The Journal of Experimental Medicine 205, 105–115 (2008). 10.1084/jem.20071367

42 Ganchua, S. K. C. et al. Lymph nodes are sites of prolonged bacterial persistence during Mycobacterium tuberculosis infection in macaques. PLOS Pathogens 14, e1007337 (2018). 10.1371/journal.ppat.1007337

43 Ganchua, S. K. C., White, A. G., Klein, E. C. & Flynn, J. L. Lymph nodes—The neglected battlefield in tuberculosis. PLOS Pathogens 16, e1008632 (2020). 10.1371/journal.ppat.1008632

44 Zhang, J. et al. Erdman infection of cynomolgus macaques of Chinese origin. J Thorac Dis 10, 3609–3621 (2018). 10.21037/jtd.2018.05.189

45 Hansen, S. G. et al. Prevention of tuberculosis in rhesus macaques by a cytomegalovirus-based vaccine. Nature Medicine 24, 130–143 (2018). 10.1038/nm.4473

46 Rayner, E. L. et al. Early lesions following aerosol infection of rhesus macaques (Macaca mulatta) with Mycobacterium tuberculosis strain H37RV. J Comp Pathol 149, 475–485 (2013). 10.1016/j.jcpa.2013.05.005

47 Choreño-Parra, J. A. et al. Mycobacterium tuberculosis HN878 Infection Induces Human-Like B-Cell Follicles in Mice. The Journal of Infectious Diseases 221, 1636–1646 (2020). 10.1093/infdis/jiz663

48 Cohen, S. B. et al. Host and pathogen genetic diversity shape vaccine-mediated protection to Mycobacterium tuberculosis. Frontiers in Immunology 15 (2024). 10.3389/fimmu.2024.1427846

49 Bucşan, A. N. et al. Response to Hypoxia and the Ensuing Dysregulation of Inflammation Impacts. Am J Respir Crit Care Med 206, 94–104 (2022). 10.1164/rccm.202112-2747OC

50 Bishai, W. R. et al. Virulence of *Mycobacterium tuberculosis* CDC1551 and H37Rv in Rabbits Evaluated by Lurie’s Pulmonary Tubercle Count Method. Infection and Immunity 67, 4931–4934 (1999). 10.1128/iai.67.9.4931-4934.1999

