## Supplemental Figure for "Protection against reinfection with *Mycobacterium tuberculosis* extends across heterologous Mtb lineages"

Supplemental Figures and Legends

**Supplemental Figure S1**

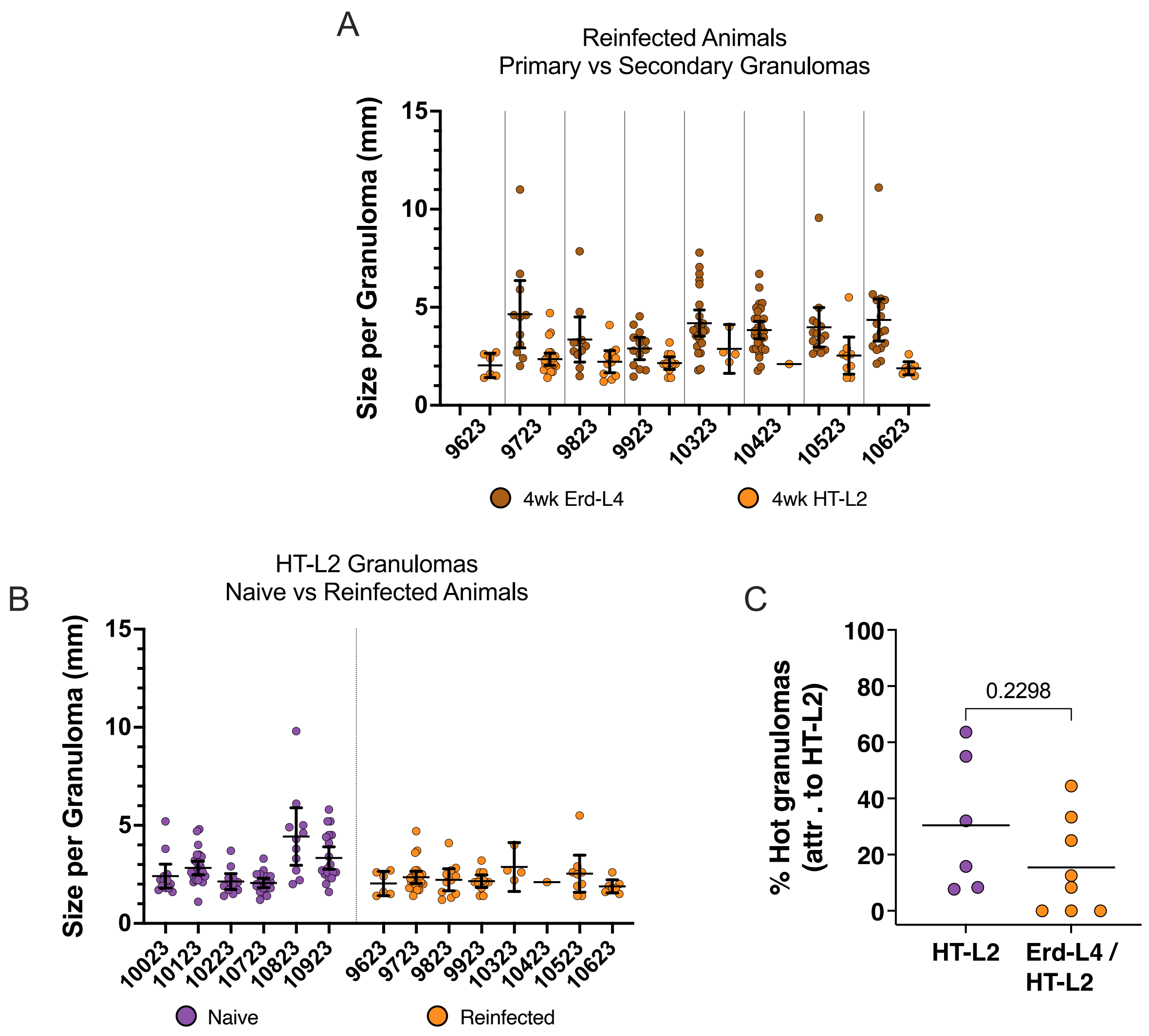


**Supplemental Figure S1. Serial PET CT analysis of granulomas.** (**A**) Size of granulomas that appeared 4 weeks post-primary (dark orange) and post-secondary (light orange), measured by PET CT scans. 9623 had large consolidations by 4 weeks after Erd-L4 challenge, so no individual granulomas are reported for that time point. (**B**) Size of granulomas at necropsy following HT-L2 challenge for naïve (purple) and reinfected (orange) animals. Points shown for the reinfection group are the same shown in panel A (light orange). For panels A and B, error bars represent mean and 95% confidence interval. (**C**) Percentage of HT-L2 granulomas per animal that were "hot" by PET CT scans, determined by a peak SUV greater than 2.3. In panel C, each symbol represents and animal, lines represent the group median, and p values are reported for Welch's unpaired t tests.

**Supplemental Figure S2**

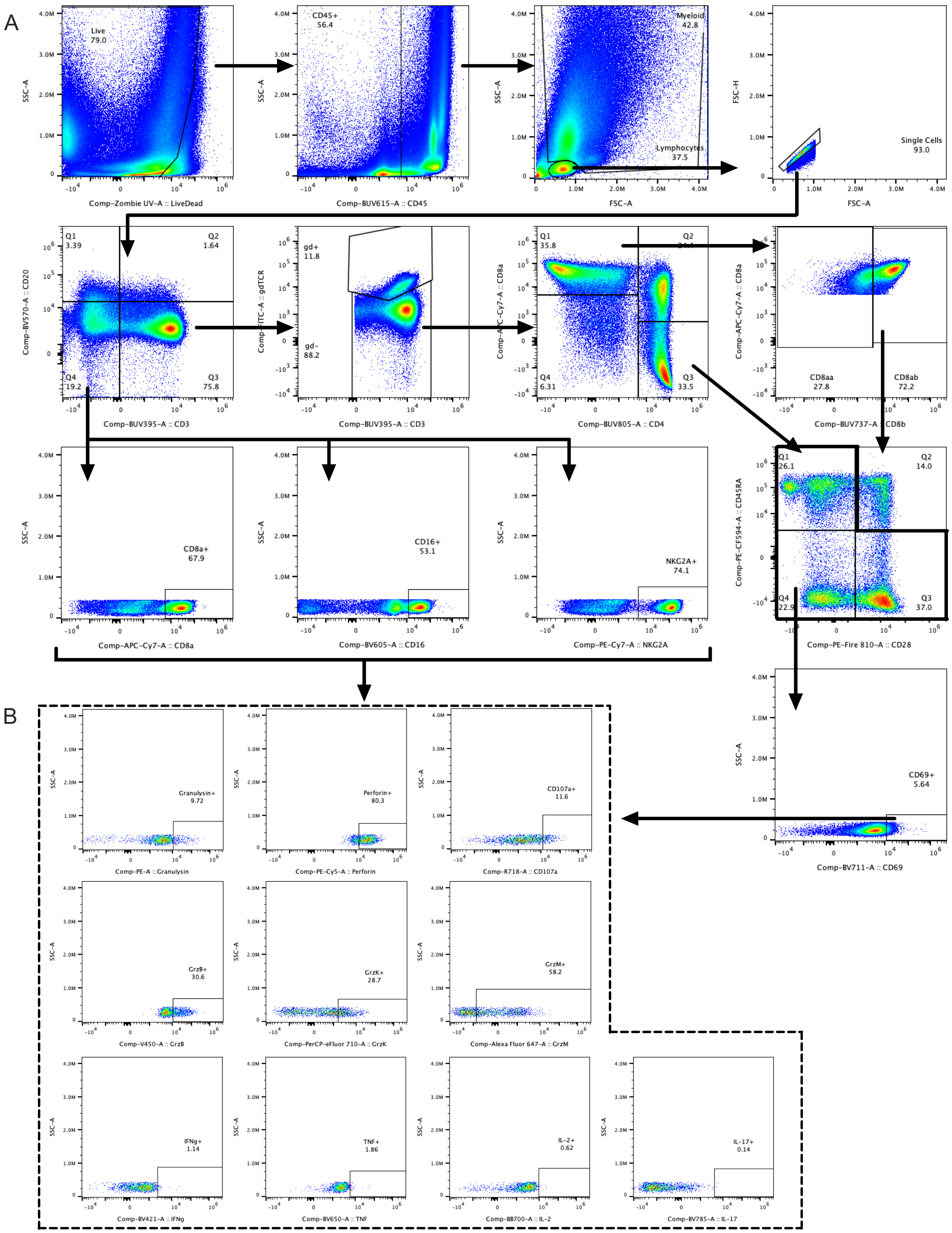


**Supplemental Figure S2. Representative flow cytometry gating strategy.** (**A,B**) Identification of cell phenotype (A) and functional profile (B).
